# Atenolol reduces cardiac-mediated mortality in genetic mouse model of sudden unexpected death in epilepsy

**DOI:** 10.1101/2023.12.10.570964

**Authors:** Ming S. Soh, Alibek Kuanyshbek, Erlina S. Mohamed Syazwan, Hian M. Lee, Chaseley E. McKenzie, A. Marie Phillips, Amanda Hu, Ingrid E. Scheffer, Christopher Semsarian, Samuel F. Berkovic, Christopher A. Reid

## Abstract

Sudden Unexpected Death in Epilepsy (SUDEP) is the leading cause of premature mortality in epilepsy. Genetic cardiac risk factors, including loss-of-function *KCNH2* variants, have been linked to SUDEP. We hypothesised that seizures and LQTS interact to increase SUDEP risk. To investigate this, we crossed *Kcnh2*^+/-^ and *Gabrg2*^R43Q/+^ mice that model LQTS and genetic epilepsy, respectively. Electrocorticography and electrocardiogram confirmed that *Kcnh2*^+/-^ mice had a LQTS phenotype, while *Gabrg2*^R43Q/+^ mice displayed spontaneous seizures. Double mutant mice (*Gabrg2*^R43Q/+^/*Kcnh2*^+/-^) had both seizure and LQTS phenotypes that were indistinguishable from the respective single mutant mice. Survival analysis revealed that *Gabrg2*^R43Q/+^/*Kcnh2*^+/-^ mice experienced a disproportionate higher rate of seizure-related death. Long-term oral administration of atenolol, a cardiac-selective β-blocker, significantly improved survival in the *Gabrg2*^R43Q/+^/*Kcnh2*^+/-^ mice. An additional mouse model, *Hcn1^M294L^*^/+^/*Kcnh2*^+/-^, based on a *HCN1* developmental epileptic encephalopathy variant, also experienced a disproportionately higher rate of premature death that was rescued by atenolol. *Kcnh2*^+/-^ mice also spent more time in ventricular arrhythmia during proconvulsant-induced seizures. Overall, the data implicates cardiac and loss-of-function *KCNH2* variants as an important risk factor, and the potential repurposing of β- blockers as a prevention strategy, for SUDEP in a subset of epilepsy patients.

## INTRODUCTION

Sudden unexpected death in epilepsy (SUDEP) is the most common cause of premature mortality for the estimated 50 million people with epilepsy globally^1^. The International League Against Epilepsy defines SUDEP as a sudden, unexpected death in a person with epilepsy, with or without evidence for a seizure preceding the death, in which there is no evidence of other disease, injury, or drowning that caused the death^2,3^. SUDEP often occurs among young adults and is second only to stroke in years of potential life lost among neurological diseases, making it the most catastrophic complication of epilepsy^4^. As most cases are not witnessed at the time of death, it remains a challenge to dissect out the exact cause underlying SUDEP. There are several recognised SUDEP risk factors including frequent generalised tonic-clonic seizures, poor treatment adherence, and resistance to anti-seizure medications^5–8^. Nonetheless, patients with well-controlled seizures can still experience SUDEP^7–9^. The involvement of neuro-cardio-respiratory pathways is hypothesised to be the primary causes of death in SUDEP^10–13^. The MORTEMUS study suggested a unifying ‘respiratory hypothesis’ with 67% of the studied SUDEP cases showing that terminal apnoea preceded terminal asystole^14^. However, the mechanism is more complex, as the remaining 33% of cases indicated that terminal events were consistent with cardiorespiratory arrest. Furthermore, significant cardiac events including bradycardia and transient asystole preceded terminal apnoea in most patients^14,15^. It is therefore likely that the underlying cause of SUDEP is complex, multifactorial and may vary between patients.

Cardiac arrhythmia has been regularly proposed as one of the possible causes of SUDEP^12^. One reason relates to the similarities between SUDEP and sudden cardiac death due to arrhythmias^16^. Furthermore, changes in cardiac function, including an increase in heart rate and life-threatening arrhythmia can occur during epileptic seizures^17–19^. Recently, a retrospective study demonstrated that prolonged QT interval on the electrocardiogram (ECG) could predict all-cause mortality in patients with epilepsy^20^. In two other retrospective studies, lower heart rate variability was found to be associated with a higher SUDEP risk^21,22^. With increased genetic testing in recent years, pathogenic variants in major cardiac genes such as *KCNH2* and *SCN5A*, which cause inherited arrhythmia syndromes, have been identified in up to 15% of SUDEP cases^10,23–25^. Pathogenic *KCNH2* variants, in particular, were most frequently implicated in a large genetic analysis of SUDEP patients^23^.

*KCNH2*, also known as the human Ether-à-go-go-Related Gene or hERG, encodes the potassium ion channel Kv11.1, which is expressed widely in the heart and brain^16^. In the heart, this channel is responsible for regulating the delayed rectifier potassium current, Ikr, critical for phase 3 repolarisation following a cardiac action potential. Loss-of-function pathogenic *KCNH2* variants cause a congenital arrhythmia condition known as long-QT syndrome (LQTS)^16^. One of the characteristics of LQTS is the prolongation of the QT interval on the ECG that increases the risk of a severe form of ventricular arrhythmia known as ‘torsades de pointes’, which in turn can trigger sudden cardiac death^16,26^. LQTS caused by loss-of-function *KCNH2* variants accounts for approximately 30% of all LQTS cases^16^. In the brain, Kv11.1 regulates neuronal excitability and has been associated with seizure and epilepsy^27–31^. However, as patients with LQTS often experience syncope accompanied by a non-epileptic seizure, they are frequently misdiagnosed with epilepsy and incorrectly treated with anti-seizure medications^28,32,33^.

Our recent study found a 3-fold enrichment of *KCNH2* variants that caused a loss of Kv11.1 function in individuals who experienced SUDEP compared with living epilepsy patients^34^. This suggests that loss-of-function *KCNH2* variants can increase SUDEP risk. We hypothesised that the increased risk of sudden death is due to the additive effects of seizure- and arrhythmia-mediated cardiac dysfunction. Furthermore, we postulated that repurposing a LQTS treatment could decrease the risk of SUDEP. To test this, we developed a novel SUDEP mouse model that has both genetic epilepsy and *Kcnh2*-mediated cardiac arrhythmia. Our results demonstrated that loss-of-function *Kcnh2* significantly reduced survival in a mouse model of genetic epilepsy without affecting seizure susceptibility. Importantly, oral treatment with the cardiac-selective β-blocker, atenolol, was able to rescue double mutant mice from premature death. Atenolol was also able to reduce mortality in a second genetically-based SUDEP mouse model. Further investigation on the impact of induced seizures on cardiac electrophysiology revealed an increased propensity to ventricular arrhythmia in the loss-of-function *Kcnh2* mouse model, which was reversed by atenolol. Our study provides evidence that genetic cardiac arrhythmia acts as a risk factor for SUDEP, and that the administration of β-blockers may be a useful preventative strategy in patients with epilepsy who have a higher genetic risk for SUDEP due to loss-of-function *KCNH2* variants.

## RESULTS

To develop a genetic model of SUDEP, *Gabrg2*^R43Q/+^ and *Kcnh2*^+/-^ mice were crossed to produce a double mutant model. The *Gabrg2*^R43Q/+^ mouse is a well-established model of generalised epilepsy based on a pathogenic arginine to glutamine variant at position 43 in GABAA γ2 subunit gene *GABRG2*^35–38^*. Kcnh2*^+/-^ mice have loss-of-function *KCNH2* and model LQTS^39–41^. Mating between *Gabrg2*^R43Q/+^ and *Kcnh2*^+/-^ mice produced littermates which were born in Mendelian ratio of the four genotypes (wild-type, *Gabrg2*^R43Q/+^, *Kcnh2*^+/-^, *Gabrg2*^R43Q/+^/*Kcnh*2^+/-^). All mice appeared healthy with weights of the littermates similar across age for the different genotypes (Supplemental Figure 1 and Table 1).

**Table 1.** Summary of weight, ECoG and ECG comparison between the four genotypes.

| **Body weight (g)** |  |  |  |  |
| --- | --- | --- | --- | --- |
|  | Wild-type<br>(N=45) | *Gabrg2*<sup>R43Q/+</sup><br>(N=41) | *Kcnh2*<sup>+/-</sup> (N=37) | *Gabrg2*<sup>R43Q/+</sup>/<br>*Kcnh2*<sup>+/-</sup> (N=38) |
| P23-40 | 15.59 ± 0.46 | 15.93 ± 0.68 | 16.43 ± 0.69 | 16.10 ± 0.55 |
| P40-60 | 20.07 ± 0.55 | 20.28 ± 0.61 | 21.38 ± 0.54 | 20.48 ± 0.23 |
| P60-80 | 23.17 ± 0.48 | 21.40 ± 0.70 | 22.87 ± 0.58 | 21.83 ± 0.36 |
| P80-100 | 24.22 ± 0.87 | 25.19 ± 0.99 | 26.34 ± 0.33 | 25.15 ± 0.80 |
| **Heart weight: body weight ratio (mg/g)** |  |  |  |  |
|  | Wild-type<br>(N=10) | *Gabrg2*<sup>R43Q/+</sup><br>(N=6) | *Kcnh2*<sup>+/-</sup> (N=10) | *Gabrg2*<sup>R43Q/+</sup>/<br>*Kcnh2*<sup>+/-</sup> (N=6) |
| P70-80 | 7.59 ± 0.47 | 7.71 ± 0.31 | 7.52 ± 0.28 | 7.21 ± 0.42 |
| **ECoG parameters** |  |  |  |  |
|  | Wild-type<br>(N=10) | *Gabrg2*<sup>R43Q/+</sup><br>(N=13) | *Kcnh2*<sup>+/-</sup> (N=10) | *Gabrg2*<sup>R43Q/+</sup>/<br>*Kcnh2*<sup>+/-</sup> (N=12) |
| Number of spikes per hour | 0.06 ± 0.04 | 9.42 ± 1.96**** | 0.05 ± 0.05 | 8.20 ± 1.68*** |
| Number of spike-wave discharges per hour | 0.37 ± 0.14 | 12.75 ± 3.25** | 0.47 ± 0.22 | 11.05 ± 3.87** |
| Number of seizures Racine 3 and above | 0.00 ± 0.00 | 1.08 ± 0.67 | 0.00 ± 0.00 | 1.00 ± 1.00 |
| **ECG parameters** |  |  |  |  |
|  | Wild-type (N=6) | *Gabrg2*<sup>R43Q/+</sup><br>(N=5) | *Kcnh2*<sup>+/-</sup> (N=5) | *Gabrg2*<sup>R43Q/+</sup>/<br>*Kcnh2*<sup>+/-</sup> (N=6) |
| Heart rate (beats per minute) | 548.30 ± 14.93 | 573.20 ± 11.62 | 560.00 ± 20.74 | 557.30 ± 12.62 |
| Heart rate variability (RMSSD, ms) | 4.10 ± 0.64 | 4.08 ± 0.86 | 4.03 ± 0.75 | 4.26 ± 1.06 |
| P-wave duration (ms) | 12.11 ± 0.14 | 12.14 ± 0.16 | 12.15 ± 0.17 | 11.95 ± 0.16 |
| PR interval (ms) | 35.27 ± 0.22 | 35.26 ± 0.22 | 35.7 ± 0.29 | 35.05 ± 0.25 |
| QRS interval (ms) | 13.55 ± 0.11 | 13.22 ± 0.13 | 13.40 ± 0.17 | 13.48 ± 0.13 |
| RR interval (ms) | 107.50 ± 0.94 | 108.3 ± 1.42 | 107.4 ± 1.56 | 105.6 ± 1.27 |
| QT interval (ms) | 41.38 ± 0.84 | 40.88 ± 0.37 | 46.48 ± 0.76**** | 46.44 ± 0.43**** |
| QTc interval (ms) | 39.77 ± 0.67 | 39.37 ± 0.32 | 44.42 ± 0.87**** | 44.96 ± 0.56**** |

Note: Statistical comparison was made with wild-type values. Data presented as mean ± s.e.m.
\*\* P < 0.01
\*\*\* P < 0.001
\*\*\*\* P < 0.0001

### *Kcnh2* knockout does not alter the seizure phenotype or electrocorticogram marker of the ***Gabrg2*^R43Q/+^** mouse model of generalised seizures

We first characterised the baseline epileptiform activity of littermates of the four genotypes through 24-hour video electrocorticogram (ECoG) recording (Figure 1, A–G, and Table 1). We confirmed that the *Gabrg2*^R43Q/+^ mice recapitulated generalised epilepsy features seen in humans, including interictal spikings (Figure 1, B and C) and spike and wave discharges (SWDs) (Figure 1, D and E)^35,36^. A subset of *Gabrg2*^R43Q/+^ mice also experienced spontaneous Racine 3-4 seizures with forelimb clonus, rearing and extended tail (Figure 1, F and G, and Supplemental Movie 1). Double mutant *Gabrg2*^R43Q/+^/*Kcnh*2^+/-^ mice also exhibit interictal spikings, SWDs and rare spontaneous Racine 3-4 class seizures (Figure 1, B–G). Importantly, comparison of seizure phenotype between *Gabrg2*^R43Q/+^ and *Gabrg2*^R43Q/+^/*Kcnh2*^+/-^ mice revealed no difference in the frequencies of interictal spikes, SWDs or spontaneous seizures (Figure 1, B–G). As expected, ECoG recording revealed no spontaneous seizures and only a minimal number of interictal spikes and SWDs in the wild-type and single mutant *Kcnh2*^+/-^ mice (Figure 1, A–G; Table 1). These data suggest that heterozygous knockout of *Kcnh2* does not modulate epileptiform or spontaneous seizure activity in the *Gabrg2*^R43Q/+^ mouse model.

**Figure 1.**
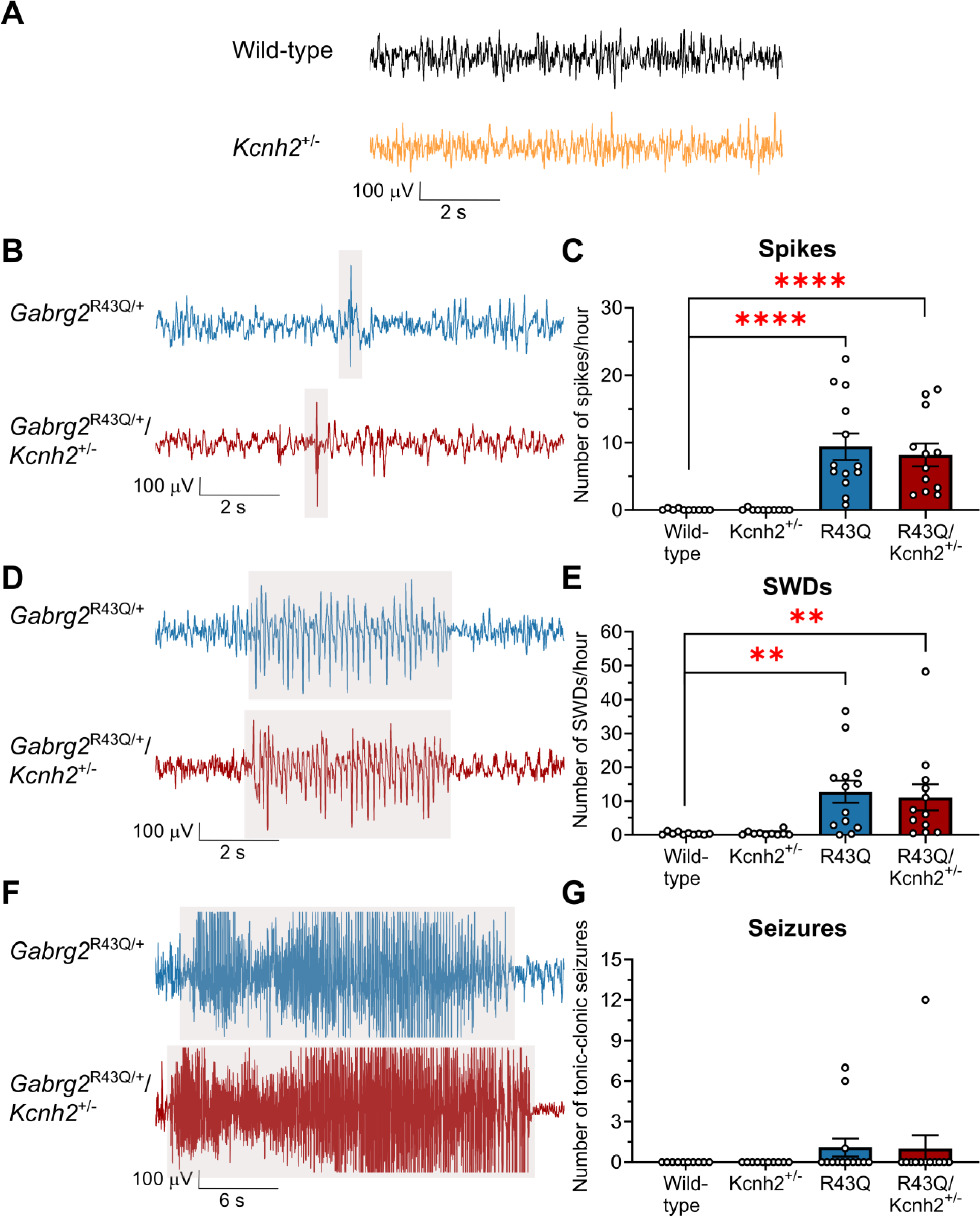
Characterisation of baseline epileptiform activity in all four genotypes of *Gabrg2*^R43Q/+^ and *Kcnh2*^+/-^ mating. **(A)** Example ECoG traces from a wild-type and a *Kcnh2*^+/-^ mouse showing normal brain electrical activity. **(B-G)** Respective grey-highlighted areas in sample ECoG traces on the left from *Gabrg2*^R43Q/+^ and *Gabrg2*^R43Q/+^/*Kcnh2*^+/-^ mice, and graphs on the right with mean ± s.e.m. comparing the four genotypes, showing **(B-C)** interictal spike occurrence per hour, **(D-E)** abnormal discharges of approximately 6-8 Hz spike-wave complex (SWDs) per hour, and **(F-G)** total number of ictal activities throughout the 24-hour recording. ** P < 0.01, **** P < 0.0001 compared to wild-type using Kruskal-Wallis test with Dunn’s posthoc, N=10-13 mice/group. See Table 1 for summary of comparison.

### Prolonged QT interval in *Kcnh2*^+/-^ and *Gabrg2*^R43Q/+^/*Kcnh2*^+/-^ mice

24-hour video-ECG recordings were made to measure cardiac parameters in littermates of the *Gabrg2*^R43Q/+^ and *Kcnh2*^+/-^ mating (Figure 2). Recordings revealed no spontaneous arrhythmias in either the single *Kcnh2*^+/-^ nor double mutant mice during this period. However, a prolonged corrected QT interval (QTc) was observed in the *Kcnh2*^+/-^ mice compared to wild-type mice, thus confirming a LQTS phenotype (Figure 2, A and B)^39–41^. QTc intervals did not differ between *Gabrg2*^R43Q/+^ and WT mice (Figure 2, A and B). *Gabrg2*^R43Q/+^/*Kcnh2*^+/-^ mice also have a prolonged QTc interval suggesting that the LQTS phenotype is maintained in the double mutant mice (Figure 2, A and B). There was no difference in the heart rate, root mean square of successive differences between normal heartbeats (RMSSD, which is a measure of heart rate variability), RR interval, P-wave duration, or PR interval between the four genotypes (Figure 2, C–G). Heart weight to body weight ratio was also similar between all groups (Table 1). All ECG parameters are presented in Table 1.

**Figure 2.**
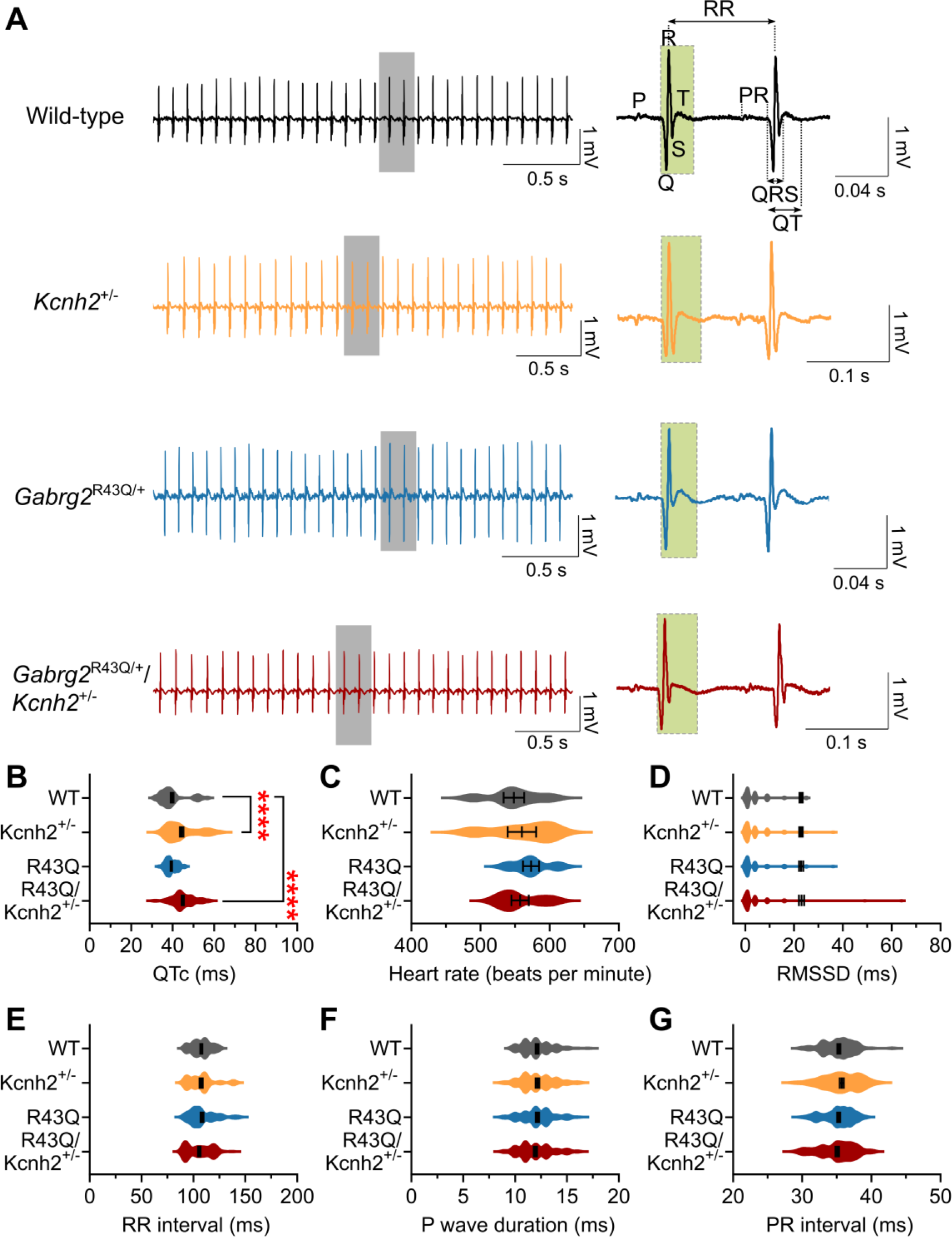
Loss-of-function *Kcnh2* causes QT prolongation in mice. **(A)** Example ECG traces showing sinus rhythm from a single wild-type, *Kcnh2*^+/-^, *Gabrg2*^R43Q/+^ or *Gabrg2*^R43Q/+^/*Kcnh2*^+/-^ mouse. The right traces, showing PQRST waves, are zoomed in from the **grey**-highlighted boxes from the respective left traces. The **green**-highlighted boxes on the zoomed-in traces on the right emphasise the difference in QT intervals between the four genotypes. **(B-G)** Violin distribution plots with mean ± s.e.m. comparing **(B)** corrected QT interval (QTc), **(C)** heart rate, **(D)** root mean square of successive differences between normal heartbeats (RMSSD), which is a measure of heart rate variability, **(E)** RR interval, **(F)** P-wave duration, and **(G)** PR interval. **** P < 0.0001 compared to wild-type using Kruskal-Wallis test with Dunn’s posthoc, N=5-6 mice/group. All ECG parameters are summarised in Table 1.

### Loss of *Kcnh2* function increases spontaneous death rate in *Gabrg2*^R43Q/+^ genetic epilepsy mice

To evaluate the impact of having both genetic epilepsy and LQTS on SUDEP risk, we performed long-term 24/7 video recordings on littermates from weaning (P21-P23) until P100. The mice were group-housed according to sex in a quiet room with a normal light-dark cycle. We scored the time of death and evaluated the associated behaviour, including seizures, during death. All wild-type mice survived until the end of the long-term recording at P100 (Figure 3). A small number of *Kcnh2*^+/-^ (2 out of 37) and *Gabrg2*^R43Q/+^ (4 out of 41) mice died prematurely during the recording period. The *Gabrg2*^R43Q/+^ and *Gabrg2*^R43Q/+^/*Kcnh2*^+/-^ mice were found dead with hind-limb extension. Video recordings confirmed that the mice experienced Racine class 4-5 seizures resulting in death. In contrast, the deceased *Kcnh2*^+/-^ single mutant mice did not exhibit hind-limb extension nor were seizures observed prior to death.

**Figure 3.**
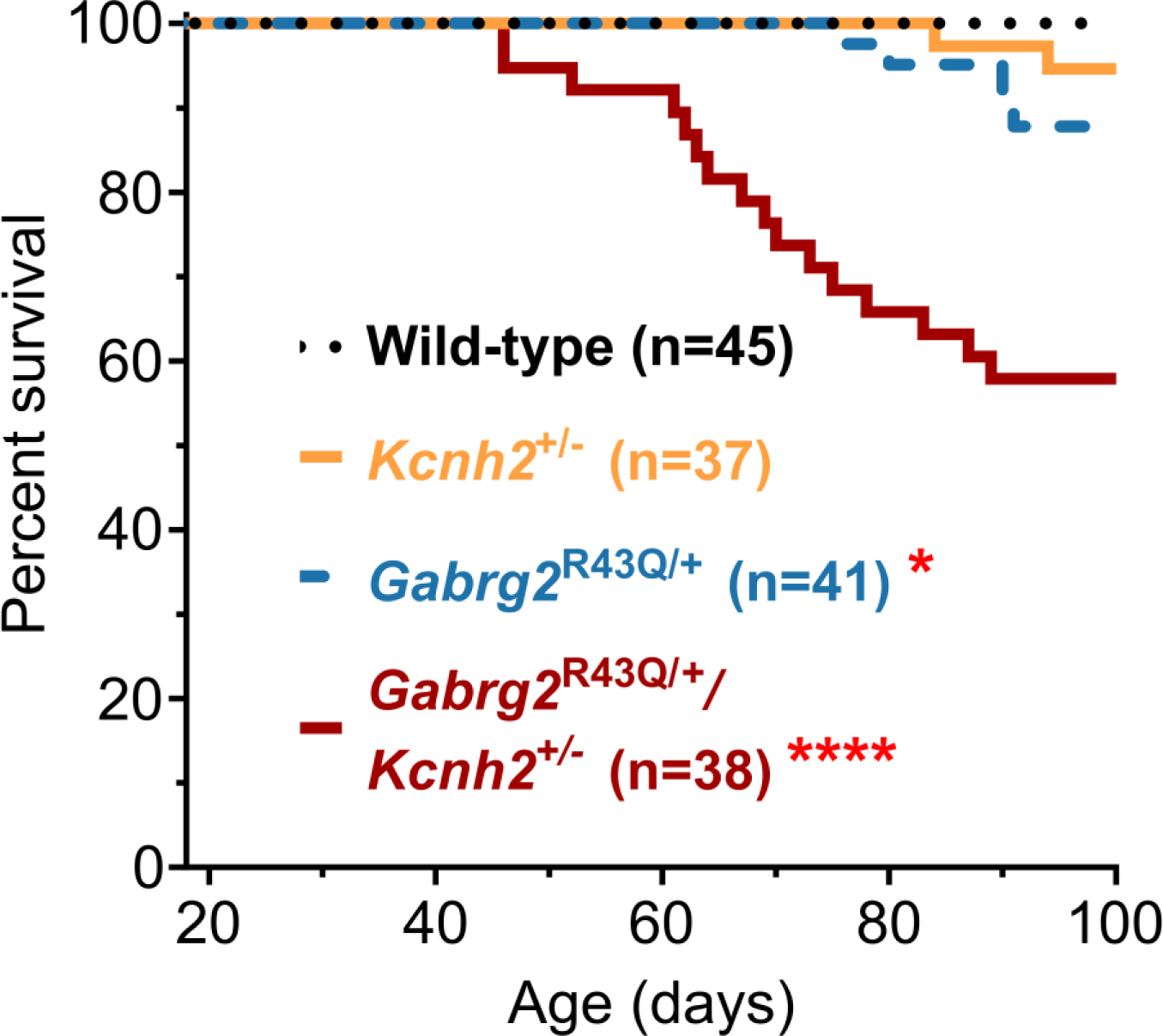
Loss of *Kcnh2* function increases risk of premature mortality in *Gabrg2*^R43Q/+^/*Kcnh2*^+/-^ SUDEP mouse model. Kaplan-Meier survival curve demonstrating a lower survival rate in the *Kcnh2*^+/-^, *Gabrg2*^R43Q/+^ and *Gabrg2*^R43Q/+^/*Kcnh2*^+/-^ mice. Video recordings confirmed that the *Gabrg2*^R43Q/+^ and *Gabrg2*^R43Q/+^/*Kcnh2*^+/-^ mice died during Racine 4-5 seizure. * P < 0.05, **** P < 0.0001 compared to wild-type using Mantel-Cox log-rank test; N=37-45 mice/group.

An increase in spontaneous death rate in the *Gabrg2*^R43Q/+^ and *Kcnh2*^+/-^ single mutant mice suggests that both pathogenic variants can independently increase the risk of premature death. Strikingly, double mutant *Gabrg2*^R43Q/+^/*Kcnh2*^+/-^ mice were disproportionately more likely to die prematurely within the timeframe (16 out of 38, Figure 3), suggesting a synergistic impact of the two independent risk factors. As *Kcnh2*^+/-^ does not worsen seizure frequency or severity, the increased in premature death in the *Gabrg2*^R43Q/+^/*Kcnh2*^+/-^ mice is likely caused by cardiac dysfunction in addition to the existing risk caused by seizures.

### Atenolol improves survival in *Gabrg2*^R43Q/+^/*Kcnh2*^+/-^ mice

β-blockers are first-line therapy for patients with LQTS caused by loss-of-function *KCNH2* variants and are effective in reducing the risk of sudden cardiac death^42–45^. We therefore hypothesised that β-blockers may reduce the risk of premature death in our preclinical model of SUDEP. Specifically, we used the cardiac-selective atenolol, which has a higher affinity for the β1-adrenergic receptor and does not cross the blood-brain barrier^46^. We randomly assigned *Gabrg2*^R43Q/+^/*Kcnh2*^+/-^ mice to either atenolol-treated or untreated group. In the atenolol-treated groups, we dissolved 0.1 or 0.6 g/L atenolol in tap drinking water^47^ and introduced the treatment from P40. The time point P40 was chosen as this was before the first death was recorded in the *Gabrg2*^R43Q/+^/*Kcnh2*^+/-^ mice (Figure 3). Untreated mice were given standard tap drinking water. The average volume drunk was similar between untreated and atenolol-treated groups (Supplemental Figure 2) and neither dose of drug caused any overt behavioural changes in the mice. While the lower atenolol dose was ineffective, the higher 0.6 g/L atenolol dose significantly improved survival in the *Gabrg2*^R43Q/+^/*Kcnh2*^+/-^ double mutant mice (Figure 4).

**Figure 4.**
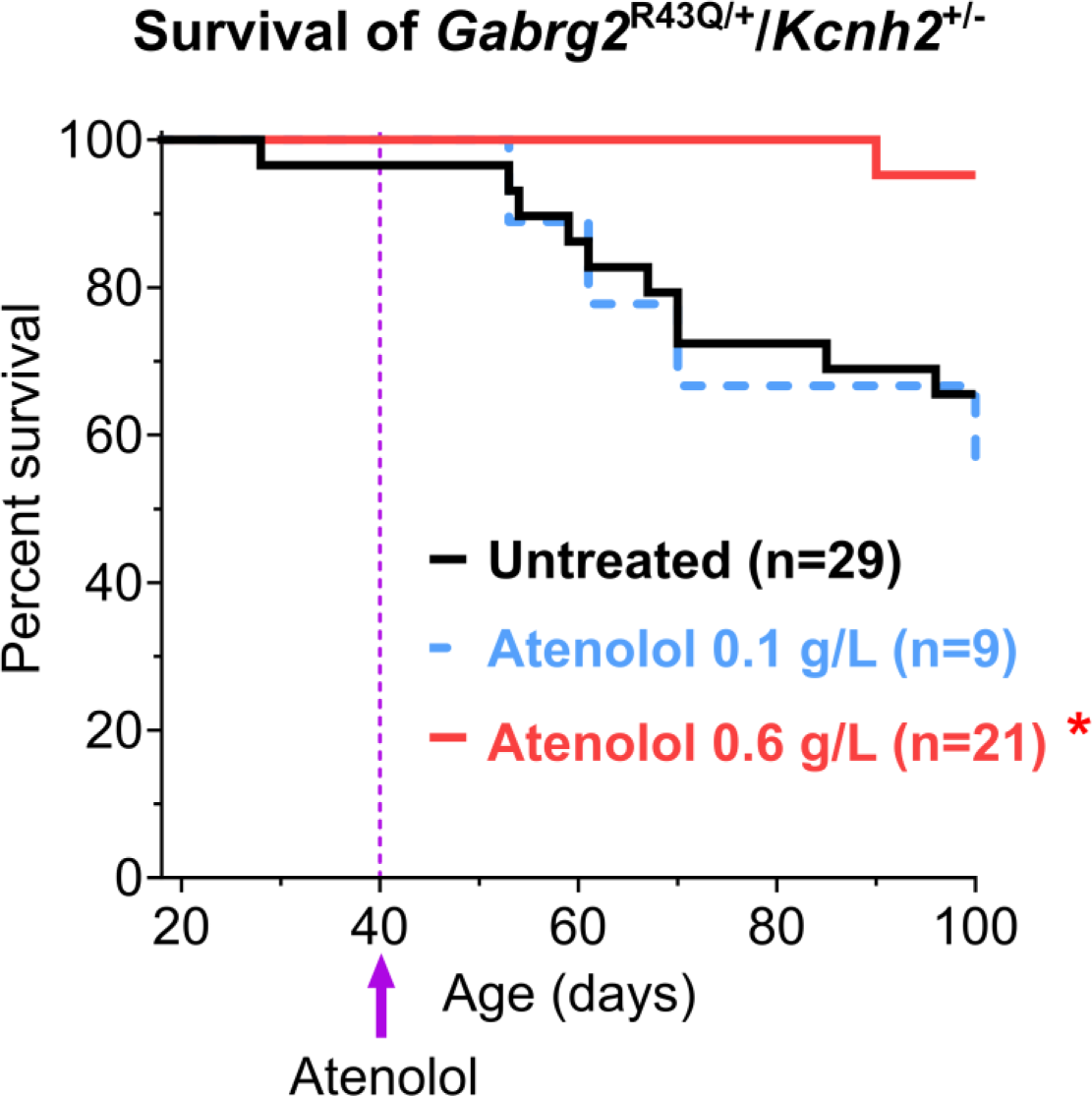
Dose-dependent β-blocker atenolol improves survival in *Gabrg2*^R43Q/+^/*Kcnh2*^+/-^ double mutant mice. **(A)** Kaplan-Meier survival curve demonstrating an improvement in survival of double mutant mice that were given 0.6 g/L atenolol, but not in mice that were given 0.1 g/L atenolol. Atenolol was introduced at P40, marked by the **purple** arrow and line. * P < 0.05 compared to untreated using Mantel-Cox log-rank test, N=9-29/group.

### Cardiac-selective impact of atenolol in *Gabrg2*^R43Q/+^/*Kcnh2*^+/-^ mice

ECG recordings performed on untreated and 0.6 g/L atenolol-treated *Gabrg2*^R43Q/+^/*Kcnh2*^+/-^ mice revealed that atenolol significantly reduced the average heart rate and QTc interval (Figure 5, A– C). Interestingly, heart rate variability RMSSD was 4.3 times higher in the atenolol-treated group (Figure 5D). Atenolol also prolonged the RR interval considerably by about 15.5 ms (Figure 5E) but had no impact on P-wave duration or PR interval (Figure 5, F and G). All ECG parameters are summarised in Table 2.

**Figure 5.**
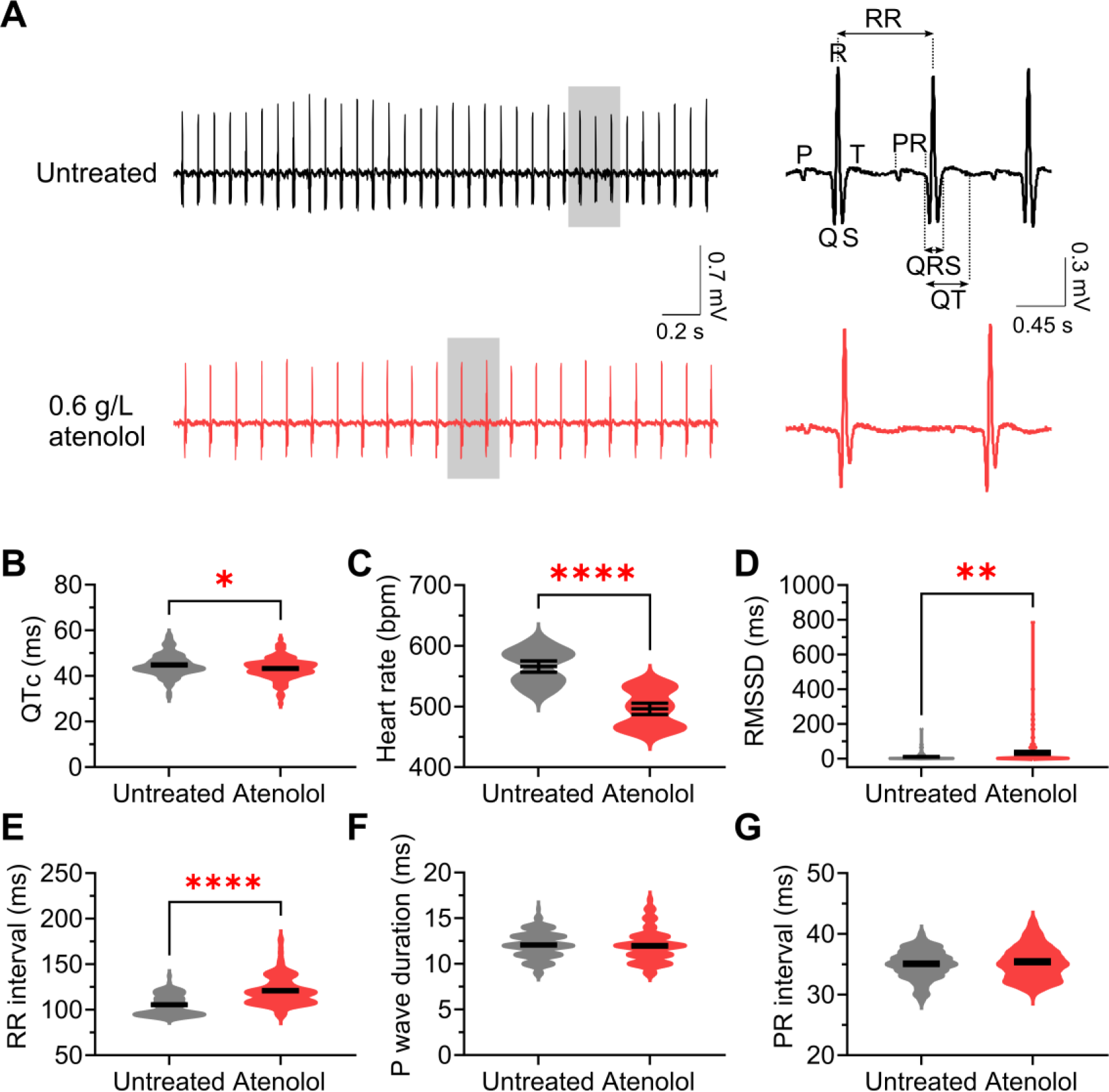
Atenolol significantly reduces heart rate and QTc in *Gabrg2*^R43Q/+^/*Kcnh2*^+/-^ mice. **(A)** Example ECG traces from a single untreated and 0.6 g/L atenolol-treated *Gabrg2*^R43Q/+^/*Kcnh2*^+/-^ mouse. The right traces are zoomed in from the **grey**-highlighted boxes from the respective left traces. **(B-G)** Violin distribution plots with mean ± s.e.m. comparing **(B)** corrected QT interval (QTc), **(C)** heart rate, **(D)** heart rate variability, **(E)** RR interval, **(F)** P-wave duration, and **(G)** PR interval. Bpm=beats per minute, * P < 0.05, ** P < 0.01, **** P < 0.0001 Mann-Whitney U-test, N=6-7 mice per group. Other ECG parameters are summarised in Table 2.

**Table 2.** Summary of ECoG and ECG comparison between untreated and atenolol-treated *Gabrg2*^R43Q/+^/*Kcnh2*^+/-^ mice.

| **ECoG parameters** |  |  |
| --- | --- | --- |
|  | Untreated (N=7) | 0.6 g/L atenolol (N=6) |
| Number of spikes per hour | 7.06 ± 2.19 | 8.23 ± 2.38 |
| Number of spike-wave discharges per hour | 9.89 ± 2.52 | 8.17 ± 2.96 |
| Number of seizures Racine 3 and above | 0.00 ± 0.00 | 0.17 ± 0.17 |
| **ECG parameters** |  |  |
|  | Untreated (N=7) | 0.6 g/L atenolol (N=6) |
| Heart rate (beats per minute) | 566.20 ± 9.39 | 496.00 ± 9.49**** |
| Heart rate variability (RMSSD, ms) | 4.93 ± 0.96 | 29.88 ± 8.52** |
| P-wave duration (ms) | 12.05 ± 0.11 | 11.95 ± 0.14 |
| PR interval (ms) | 34.98 ± 0.18 | 35.20 ± 0.22 |
| QRS interval (ms) | 13.44 ± 0.10 | 13.22 ± 0.13 |
| RR interval (ms) | 105.00 ± 0.98 | 120.50 ± 1.15**** |
| QT interval (ms) | 46.07 ± 0.35 | 47.28 ± 0.37** |
| QTc interval (ms) | 44.79 ± 0.44 | 43.20 ± 0.37* |

Note: Statistical comparison was made with wild-type values. Data presented as mean ± s.e.m.
\* P < 0.05
\*\* P < 0.01
\*\*\*\* P < 0.0001

To test if 0.6 g/L atenolol impacted seizure load, ECoG recordings were performed on untreated and atenolol-treated *Gabrg2*^R43Q/+^/*Kcnh2*^+/-^ mice (Figure 6 and Table 2). As expected, atenolol had no impact on either the interictal spiking (Figure 6, A and B) or SWD (Figure 6, C and D) in the *Gabrg2*^R43Q/+^/*Kcnh2*^+/-^ mice (Table 2). There was also no difference in the number of spontaneous seizures between both groups (Figure 6, E and F). Our results therefore suggest that atenolol lowers the risk of mortality through a cardiac-mediated mechanism.

**Figure 6.**
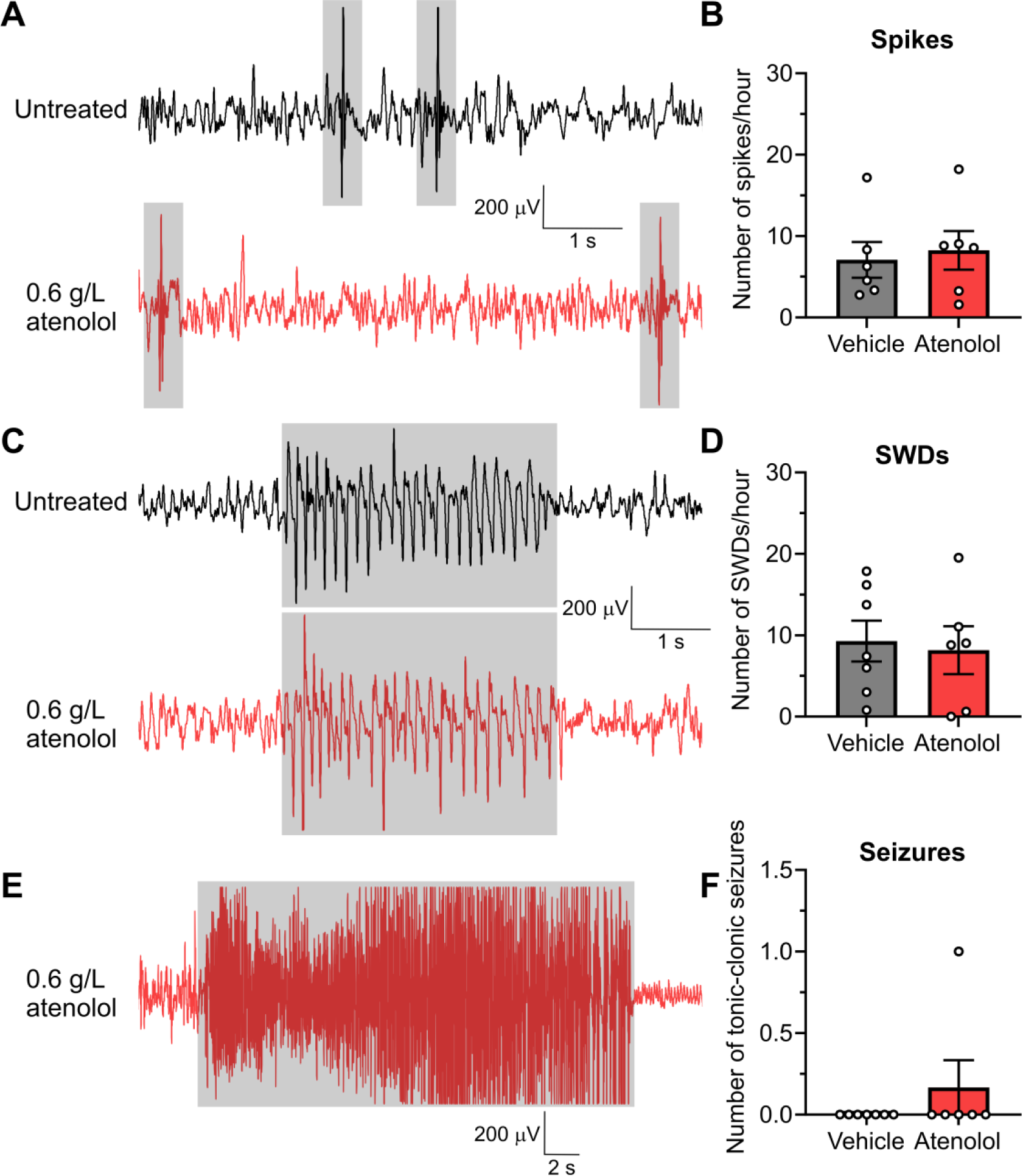
Atenolol does not affect seizure phenotype in *Gabrg2*^R43Q/+^/*Kcnh2*^+/-^ mice. **(A-F)** Respective **grey**-highlighted areas in sample ECoG traces on the left from untreated or 0.6 g/L atenolol-treated *Gabrg2*^R43Q/+^/*Kcnh2*^+/-^ mice, and graphs on the right with mean ± s.e.m. comparing between the two groups, showing **(A-B)** number of spikes per hour, **(C-D)** spike-wave discharge events (SWDs) per hour, and **(E-F)** total number of seizures within a 24-hour period (see Table 2). P > 0.05 between the two groups for all three parameters. Student’s t-test was used for **(B)** and **(D)**, while Mann-Whitney U-test was used for **(F)**, N=6-7 mice per group.

### Atenolol reduces risk of premature death in *Hcn1*^M294L/+^/*Kcnh2*^+/-^ mice

To investigate if loss of *Kcnh2* function increases the risk of premature death in a different genetic epilepsy mouse model, the *Kcnh2*^+/-^ mouse was crossed with the *Hcn1*^M294L/+^ mouse model of *HCN1* developmental epileptic encephalopathy (DEE)^48^. In this cohort, no deaths were reported for either wild-type or single *Kcnh2*^+/-^ mice for the recorded period (Figure 7A). 25% (6 out of 24) of the single mutant *Hcn1*^M294L/+^ mice died prematurely (Figure 7A). Similar to that observed for the *Gabrg2*^R43Q/+^/*Kcnh*2^+/-^ mouse model, *Hcn1*^M294L/+^/*Kcnh2*^+/-^ mice were ∼2.5-fold more likely to die prematurely when compared to the single mutant *Hcn1*^M294L/+^ mice (Figure 7A). Long-term 24/7 video recordings revealed that deaths occurred following a Racine 4-5 seizure in both the single and double mutant mice. Atenolol (0.6 g/L) significantly reduced the occurrence of premature death in the *Hcn1*^M294L/+^/*Kcnh2*^+/-^ mice (Figure 7B).

**Figure 7.**
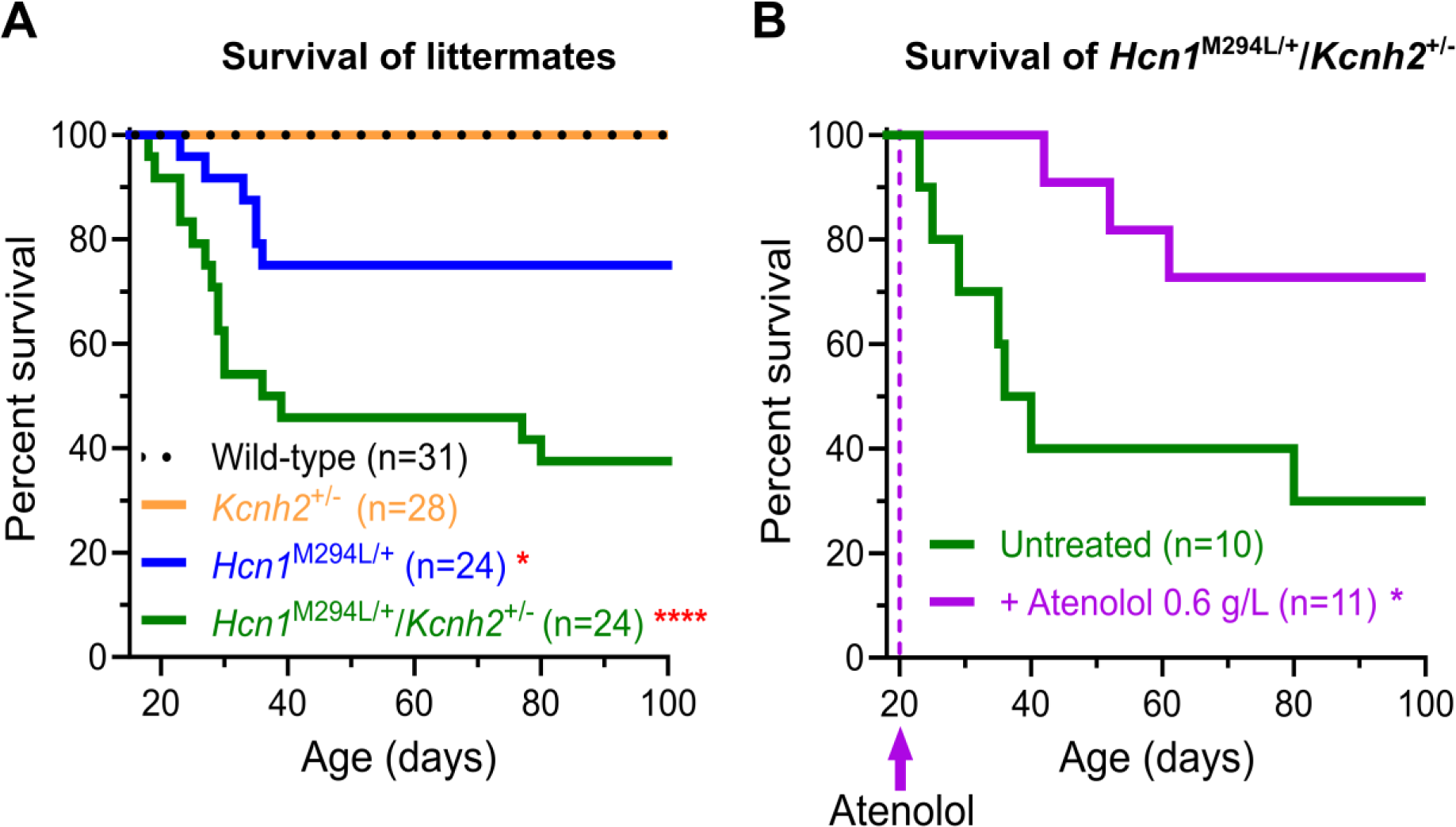
Increased risk of mortality in *Hcn1*^M294L/+^/*Kcnh2*^+/-^ SUDEP mouse model, which was rescued by cardiac-selective atenolol. **(A)** Kaplan-Meier survival curve demonstrating that *Hcn1*^M294L/+^ mice experienced higher mortality rate. However, survival is significantly reduced in the double mutant *Hcn1*^M294L/+^/*Kcnh2*^+/-^ mice. Video recordings confirmed that the mice died during Racine 4-5 seizure. N=24-31 mice/group. **(B)** Like the *Gabrg2*^R43Q/+^/*Kcnh2*^+/-^ model, cardiac-selective atenolol (0.6 g/L) was able to reduce mortality in the *Hcn1*^M294L/+^/*Kcnh2*^+/-^ mice. Atenolol was introduced at P40, marked by the **purple** arrow and line. N=10-11 mice/group. * P < 0.05, **** P < 0.0001 compared to wild-type or untreated using Mantel-Cox log-rank test.

### Loss of *Kcnh2* function increases the duration of seizure-induced ventricular arrhythmia which was rescued by atenolol

Loss-of-function *KCNH2* variants cause LQTS2 that heightens the risk of life-threatening ventricular arrhythmias in the presence of a certain physiological stressors (physical or psychological)^16,26^. We therefore hypothesise that the increase in premature death seen in both double mutant mouse models is due to an increased propensity to seizure-induced ventricular arrhythmias. However, due to the rarity and unpredictability of the seizures in both models and the technical challenge of recording ECG during spontaneous seizures causing death (caused by electromyography ‘noise’), we were unable to capture reliable cardiac arrhythmia events. To overcome this, we induced non-lethal seizures in wild-type and *Kcnh*2^+/-^mice using the proconvulsant pentylenetetrazole (PTZ). This enabled us to capture the ECG during the induction of controlled Racine 3-4 seizures and record heart function including arrhythmic events (Figure 8A). PTZ-induced seizure duration was similar between wild-type and *Kcnh*2^+/-^mice, further suggesting that seizure susceptibility was not altered in the *Kcnh*2^+/-^mice (Figure 8B, Table 3). PTZ-induced seizures increased both QTc interval and heart rate in wild-type (Table 4) and *Kcnh*2^+/-^mice (Figure 8, C and D) suggesting an increase in sympathetic drive. Importantly, *Kcnh*2^+/-^mice spent a significantly longer time in ventricular fibrillation during PTZ-induced seizures when compared to wild-type mice (Figure 8E). Administration of atenolol (10 mg/kg, i.p) was able to reduce the QTc, heart rate and time spent in ventricular fibrillation during seizure without affecting seizure susceptibility in the *Kcnh*2^+/-^mice (Figure 8, A-D; Table 3-4). The acute dose of atenolol reduced the heart rate to a similar level to that recorded for oral dosing (Figure 5C).

**Figure 8.**
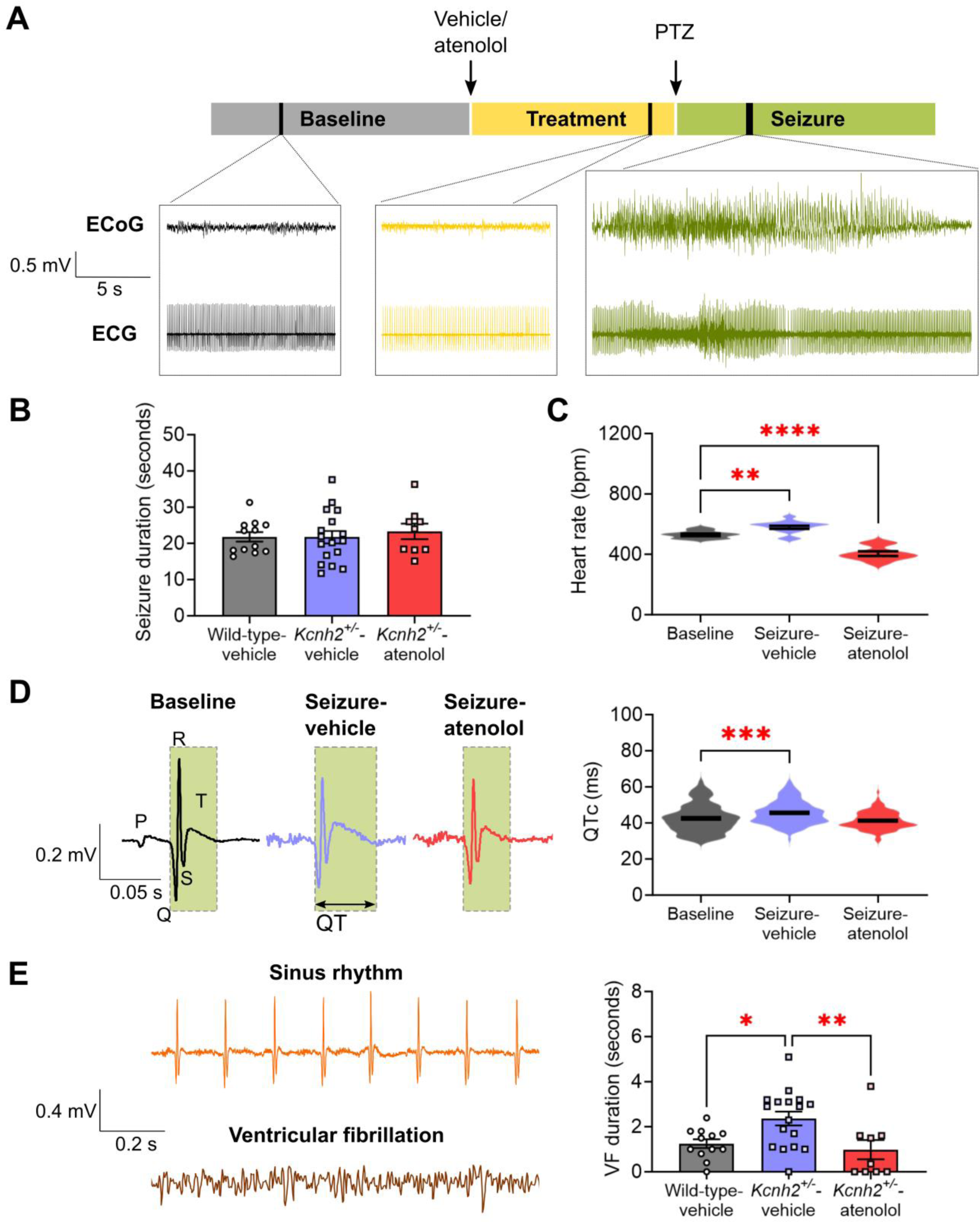
Proconvulsant-induced seizure triggers arrhythmia in wild-type and *Kcnh2*^+/-^ mice. **(A)** Experimental paradigm of proconvulsant pentylenetetrazole (PTZ)-induced seizure assay with respective sample traces of ECoG and ECG in an atenolol-treated *Kcnh2*^+/-^ mouse. **(B)** Comparison of seizure duration between wild-type and *Kcnh2*^+/-^ mice treated with vehicle (0.9% saline) or atenolol. **(C)** Impact of vehicle or atenolol treatment on heart rate in the *Kcnh2*^+/-^ mice. Bpm=beats per minute. **(D)** Left panel shows sample ECG trace recordings in *Kcnh2*^+/-^ mice, in which seizure increases QTc interval (green-highlighted boxes) in vehicle-treated mouse. Violin distribution plots with mean ± s.e.m. on the right panel summarises the QTc interval comparison. **(E)** ECG trace recordings on the left panel are respective examples of a normal sinus rhythm and ventricular fibrillation (VF) during a seizure. Comparison of VF duration between wild-type and *Kcnh2*^+/-^ mice in the presence of vehicle or atenolol is shown in the violin distribution plots with mean ± s.e.m. on the right panel. N=9-26 mice per group. * P < 0.05, ** P < 0.01, *** P < 0.001 **** P < 0.0001 One-way ANOVA with Dunnett’s post-hoc. ECG parameters are also summarised in Table 3 and 4.

**Table 3.** Comparison of seizure and ventricular fibrillation duration in vehicle-treated or 0.6 g/L atenolol-treated wild-type or *Kcnh2*^+/-^ mice induced with proconvulsant PTZ.

|  | Wild-type vehicle<br>(N=12) | *Kcnh2*<sup>+/-</sup> vehicle<br>(N=17) | *Kcnh2*<sup>+/-</sup> 0.6 g/L<br>atenolol (N=9) |
| --- | --- | --- | --- |
| Seizure duration<br>(seconds) | 21.78 ± 1.32 | 21.71 ± 1.75 | 23.31 ± 2.19 |
| Ventricular<br>fibrillation duration<br>(seconds) | 1.25 ± 0.19* | 2.37 ± 0.31 | 0.98 ± 0.42* |

Note: Statistical comparison was made with *Kcnh2*<sup>+/-</sup> vehicle values. Data presented as mean ± s.e.m.
\* P < 0.05
\*\* P < 0.01
\*\*\* P < 0.001
\*\*\*\* P < 0.00

**Table 4.** Comparison of QTc interval and heart rate in vehicle-treated or 0.6 g/L atenolol-treated wild-type or *Kcnh2*^+/-^ mice induced with proconvulsant PTZ.

| **Wild-type** |  |  |  |
| --- | --- | --- | --- |
|  | Baseline (N=6) | Vehicle (N=3) | 0.6 g/L atenolol (N=3) |
| QTc interval (ms) | 39.21 ± 0.54 | 41.55 ± 0.55** | 39.16 ± 0.46 |
| Heart rate (beats per minute) | 544.2 ± 12.21 | 590.0 ± 12.37* | 433.8 ± 23.04**** |
| ***Kcnh2*<sup>+/-</sup>** |  |  |  |
|  | Baseline (N=26) | Vehicle (N=17) | 0.6 g/L atenolol (N=9) |
| QTc interval (ms) | 42.62 ± 0.63 | 45.73 ± 0.55*** | 41.35 ± 0.46 |
| Heart rate (beats per minute) | 530.5 ± 7.76 | 580.7 ± 10.07** | 405.1 ± 15.20**** |

Note: Statistical comparison was made with baseline values. Data presented as mean ± s.e.m.
\* P < 0.05
\*\* P < 0.01
\*\*\* P < 0.001
\*\*\*\* P < 0.00

## DISCUSSION

SUDEP is a rare but catastrophic potential outcome for patients with epilepsy with no clear preventative strategy except improved seizure control. In this study, we show that loss of *Kcnh2* function increases the risk of premature death in a genetic mouse model of generalised epilepsy and a genetic mouse model of *HCN1* DEE. This is consistent with our previously published human data which demonstrated a 3-fold enrichment of loss-of-function *KCNH2* variants in SUDEP patients compared to a control group living with epilepsy^34^. Not all SUDEP mice died and the age of death spans from P45-90. This heterogeneity in survival is consistent with other animal models of SUDEP^48–50^ and reflects what human patients experience, whereby SUDEP risk is influenced by multiple factors including seizure frequency and age^5–8^. Furthermore, we demonstrate that atenolol, a cardiac-selective β-blocker, protected at-risk double mutant mice from premature death. Collectively, these data support that patients with epilepsy carrying loss-of-function *KCNH2* variants are at higher risk of SUDEP, and that preventative treatment may be effective in this cohort. Our findings will motivate future clinical studies that are needed to test this hypothesis.

While loss of *Kcnh2* function increases the risk of premature death, it does not alter spontaneous seizure susceptibility or epileptiform activity in the *Gabrg2*^R43Q/+^/*Kcnh*2^+/-^ mouse model of SUDEP. Furthermore, atenolol is effective at reducing premature death in the *Gabrg2*^R43Q/+^/ *Kcnh*2^+/-^ model without influencing seizure susceptibility. This is not unexpected, as atenolol is β1 selective and thus cardiac-specific and does not cross the blood-brain barrier^46^, which is not compromised in the *Gabrg2*^R43Q/+^ mouse model^51^. Atenolol also reduces the duration of ventricular arrhythmia caused by proconvulsant -induced seizures, without influencing seizure severity. These data suggest that pharmacological suppression of seizure-mediated sympathetic drive on the heart is sufficient to protect our SUDEP mouse model from fatal arrhythmias.

Previous studies have been conflicting as to whether dominant changes in sympathetic or parasympathetic tone result in cardiac dysfunction in patients with epilepsy^12,52–54^. However, most studies agree that tonic-clonic seizures commonly lead to a sudden increase in sympathetic response including elevated heart rate, blood pressure and changes in heart rhythm, and occasionally life-threatening ventricular fibrillation^17,55,56^. Importantly, a rise in sympathetic drive and subsequent arrhythmias caused by external stress factors are well-established triggers of sudden cardiac death in LQTS patients^57^. The increased sympathetic drive seen during a tonic-clonic seizure is therefore well positioned to act as a potential trigger of life-threatening ventricular fibrillation in at-risk individuals. Consistent with this, Van der Linde and colleagues demonstrated an increased risk of torsades de pointes due to proconvulsant-induced seizures that occurred only in dogs with drug-induced LQTS^58^. Taken together, these data suggest a logical pathological mechanism, whereby the additive effects of sympathetic ‘overdrive’ in combination with a predisposition to genetic arrhythmia may trigger an at-risk individual into life-threatening arrhythmia during a seizure.

Pathogenic variants of some arrhythmogenic genes that are expressed in both the heart and brain, such as *HCNs*, *KCNH2* and *KCNQ1*, have been suggested to cause epilepsy^27–31,52^. These variants provide an inherent risk for arrhythmia. For example, *Kcnq1* mouse models were found to experience seizures and tachyarrhythmias^52^, while *Hcn2*-knockout mice had sinus dysrhythmias^59^. However, with congenital LQTS having a prevalence of approximately 1 in 2000 people^60^, and epilepsy affecting 1% of the population worldwide^1^, a subset of patient will have epilepsy from a different cause, in addition to an existing genetic arrhythmia condition^23,25^. Findings from this study demonstrate that variants in epilepsy- and arrhythmia-causing genes can independently impact brain excitability and heart function and are therefore positioned to cause additive SUDEP risk. The observation that loss-of-function *Kcnh2* can impact survival in two quite distinct genetic causes of epilepsy suggests that our findings are generalisable and could potentially extend to other causes of epilepsy that do not have a genetic origin.

Our data also implicates arrhythmias in SUDEP more broadly. Consistent with this, rare variants in *SCN5A*, predominantly expressed in heart, that alter the biophysical properties of the encoded cardiac voltage-dependent sodium channel are found in SUDEP cases^24,61^. An array of different arrhythmia-causing genes should therefore be considered in future SUDEP studies^10,23^. Moreover, QT-prolonging medications commonly used by patients for epilepsy and associated co-morbidities, including certain antiseizure medications, antidepressants, antipsychotics, antibiotics and stimulants, could confer increased SUDEP risk^62^. Seizures are also known to indirectly change cardiac electrophysiology by altering the expression of cardiac ion channels, which subsequently could increase the risk of fatal arrhythmia^17,54,63^. A reduced heart rate variability has also been shown to associate with higher SUDEP risk^21,22^. Considering the potential role of cardiac dysfunction as a cause of SUDEP, a 12-lead ECG during initial assessment may be warranted to identify prolonged QT interval and reduced heart rate variability as potential biomarkers of increased risk. Furthermore, improved access to genetic sequencing for epilepsy patients will identify individuals with variants in arrhythmia genes that could act as biomarkers of increased SUDEP risk. Individualised therapeutic strategies are likely to be needed for different causes and further clinical investigations are essential to determine the contribution of other genetic and acquired arrhythmia factors to SUDEP risk.

Wider implementation of genetic screening in patients with epilepsy has provided an opportunity to identify likely pathogenic variants in genes causing inherited arrhythmia conditions. However, ascribing a risk of SUDEP to a given variant in an individual remains challenging. While there is a strong case of increased SUDEP risk in epilepsy patients with validated rare *KCNH2* pathogenic variants, it is less clear for more common variants that have a functional impact. For example, the *KCNH2* p.R1047L variant is found in about 3% of the population and our *in vitro* functional data revealed a small reduction in currents generated by the variant protein^34^. As such, the p.R1047L variant may be less likely to increase SUDEP risk in an individual when compared to variants that cause a larger impact on channel function. Nonetheless, at the population level, the impact can be significant given the variant’s prevalence, as it could contribute to the risk to a small degree in a large number of people. As such, an efficient and robust method is needed to ascribe functional impact of different *KCNH2* variants found in patients with epilepsy. To this end, high-throughput patch-clamp electrophysiology assay has provided evidence for the clinical interpretation of clinical *KCNH2* variants^64^. Further retrospective and prospective clinical studies are needed to determine if it will be possible to identify epilepsy patients with *KCNH2* variants who are at higher risk of SUDEP for an early intervention.

Our study is not without its limitations. The physiology of a mouse heart is significantly different to that of humans^65^. Recapitulating our findings in larger animal models and in the clinical setting will be important next steps. Furthermore, we have not specifically monitored respiratory changes in our SUDEP mice. Although the cardiac-selective β-blocker is sufficient to protect against premature death, it is possible that respiratory failure could be part of the pathogenic cascade, which warrants further study. More preclinical studies are also needed to investigate the interaction between seizures and inherited arrhythmia syndromes caused by other genes. Such preclinical models will be integral in helping to identify personalised therapeutic approaches for patients with epilepsy. For example, epilepsy patients carrying pathogenic variants in *SCN5A* may respond best to LQTS treatments that work on voltage-dependent sodium channels.

In summary, we have provided evidence that *KCNH2*-caused LQTS contributes to higher SUDEP risk in a genetic epilepsy mouse model, and that β-blockers are an effective treatment to improve survival. The genetic SUDEP mouse model will be a useful preclinical tool to further dissect underlying mechanisms and test other cardiac-targeted therapies to reduce sudden death occurrence. While a ‘unified’ pathological mechanism underlying SUDEP is unlikely, our data support the hypothesis that a cardiac-based mechanism underlies death in a subset of SUDEP.

## METHODS

### Animal study approval

All experiments involving animals were performed in accordance with the Prevention of Cruelty to Animals Act, 1986 under the guidelines of the National Health and Medical Research Council (NHMRC) of Australia Code of Practice for the Care and Use of Animals for Experimental Purposes. All procedures and monitoring include standard operating protocols that were approved by the Animal Ethics Committee at the Florey Institute of Neuroscience and Mental Health. At the conclusion of experimentation, mice were culled by cervical dislocation or decapitation following deep isoflurane anaesthesia, which are Australian & New Zealand Council for the Care of Animals in Research and Teaching (ANZCCART) approved methods.

### Animal housing and maintenance

Mice were toe-clipped and genotyped at P7, weaned at P21-23, and group-housed once weaned until experiments. The housing was within standard 15 × 30 × 12 cm cages (Techniplast, Westchester, PA, USA) maintained under 12-h dark and light (<50 lux) cycles, in ambient temperature, quiet condition, and with access to dry pellet food and tap water *ad libitum*. For each experimental comparison, age-matched littermate controls were used. Mice aged P≥25 were used for experiments. Prior to experiments, mice were acclimatised to experimental rooms for at least an hour. Sample sizes were estimated using power analysis (G*Power) based on observed effect size and a type one error rate of 0.05. Approximately equal numbers of male and female mice were used with common controls where appropriate. Blinding and randomisation of treatment and analyses (video, ECoG, ECG) to genotypes were routine.

### Animal model

Generation of the single mutant, heterozygous *Gabrg2*^R43Q/+^, *Hcn1*^M294L/+^ and *Kcnh2*^+/-^ mice were previously outlined^36,39,41,48^. *Gabrg2*^R43Q/+^ and *Kcnh2*^+/-^ strains were maintained on a DBA/2J background, *Hcn1*^M294L/+^ was maintained on a C57BL/6J. All strains had been backcrossed for at least 10 generations. The mating between *Gabrg2*^R43Q/+^ and *Kcnh2*^+/-^, and *Hcn1*^M294L/+^ and *Kcnh2*^+/-^ mice was performed on-site at the Florey Institute of Neuroscience and Mental Health, yielding wild-type, *Gabrg2*^R43Q/+^, *Kcnh2*^+/-^, or *Gabrg2*^R43Q/+^/*Kcnh*2^+/-^ offspring for the *Gabrg2*^R43Q/+^ and *Kcnh2*^+/-^ cross, and wild-type, *Hcn1*^M294L/+^, *Kcnh2*^+/-^, or *Hcn1*^M294L/+^*Kcnh*2^+/-^ offspring for the *Hcn1*^M294L/+^ and *Kcnh2*^+/-^ cross.

### Survival recording and treatment study

Once weaned, the mice were brought to an experimental room where they were weighed and group-housed according to sex and genotype. Video recording (SANNCE, Rowland Heights, CA, USA) then commenced until P100. Mice were checked daily and weighed weekly during the duration of recording, with free continuous access to food and water. Video analyses of seizures and deaths were completed off-line.

For treatment study, only the double mutant mice (*Gabrg2*^R43Q/+^/*Kcnh2*^+/-^, *Hcn1*^M294L/+^/*Kcnh2*^+/-^) were video-recorded according to the same protocol as above. At P40, mice were randomly given normal or atenolol-dissolved (Cayman Chemical, Ann Arbor, MI, USA) tap drinking water. Amount drunk was measured daily for both untreated and atenolol-treated mice. Drinking water for both groups was replaced with fresh solution every 3 days.

### Implantation electrocorticography and electrocardiogram electrodes

Instrumentation of ECoG and ECG electrodes were performed on mice aged >P25 as previously described^48^. 4-5% and 1-1.5% isoflurane was used for anaesthesia induction and maintenance, respectively. Anaesthetised mice were secured on a stereotaxic frame with heating pad. Following administration of subcutaneous anaesthetic lignocaine (6 mg/kg) and analgesic meloxicam (3 mg/kg) at the incision site perioperatively, an incision was made on the scalp to expose the skull between lambda and bregma, where three holes, each approximately 1 mm in diameter, were drilled. Two of these were positioned bilaterally over the somatosensory cortex, with the third located caudal to the lambdoid suture 0.5 mm lateral from the midline towards the right side of the skull. Stainless steel 0.10” screws (Pinnacle Technology Inc., Lawrence, KS, USA) pre-soldered with silver wire electrodes (AD instruments, Colorado Springs, CO, USA) were implanted into each hole, with the anterior two screws implanted epidurally and used as the active channel electrodes, and the posterior screw implanted slightly more shallowly and used as the reference channel electrode. A ground electrode made of silver wire was affixed to the skull immediately caudal to the lambdoid suture 0.5 mm lateral from the midline towards the left side of the skull. The silver wires of the reference, ground, and two active channel electrodes were soldered to a head mount (Pinnacle technology Inc., Lawrence, KS, USA), held in place by self-curing acrylic resin.

To place the ECG wires (Axon Connect Pte Ltd, Singapore) that had been pre-soldered to the head mount with the ECoG electrodes, an incision was made on the skin just below the head mount and on both lateral sides. Each ECG wire was tunnelled underneath the skin through the incisions, before it was sutured to the left or right muscle layer under the skin between the heart and vertebrochondral ribs with 5-0 silk suture. Incision wounds on the skin were sutured with absorbable 4-0 suture. Mice were allowed to recover for at least one week prior to experimentation.

### Electrocorticography and electrocardiogram recording

All ECoG and ECG recordings were performed on conscious mice. Each mouse was placed into individual clear plexiglass containers (140 cm x 175cm x 140 cm) with standard bedding, food and water, during each ECoG and ECG recording. The head mount was connected to a preamplifier with 10x amplification (Pinnacle Technology Inc., Lawrence, KS, USA), which in turn was connected to a data acquisition system (Sirenia 2.1.0, Pinnacle technology Inc., Lawrence, KS, USA). ECoG and ECG data were sampled at 1 kHz and 500 Hz low-pass filtered. Each set-up was also connected to an in-built dome camera (320 x 240, 20 fps, Pinnacle technology Inc., Lawrence, KS, USA) for synchronised video recording.

### Baseline epileptiform analysis

The entire 24-hour recording was visually analysed for spikes, SWDs and seizures using Sirenia software (version 2.1.0, Pinnacle technology Inc., Lawrence, KS, USA). A spike was defined by biphasic events lasting <200 ms with approximately thrice the amplitude of baseline ECoG activity^48^. SWD was characterised by burst frequency typically between 5-8 Hz, 2-3x larger amplitude than baseline, with limited spike-to-spike variability^36^. A full Racine stage 3 and above seizure generally had a combination of large-amplitude polyspike bursts and spike-wave discharges accompanied by behavioural changes such as forelimb clonus, rearing or wild jumping^66^.

### Analysis of cardiac parameters

ECG parameters were exported using Sirenia software. 30-50 consecutive beats from six sections throughout the recording were manually analysed and values averaged. For baseline comparison, only segments with reliable sinus rhythm were used for analyses. ECG parameters that were analysed included P-wave, PR, RR QRS, and QT intervals. QTc interval was calculated using the Mitchell’s formula, based on the Bazett’s formula modified for mice; 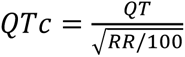 ^67,68^.

HRV was calculated based on Root Mean Square of the Successive Differences (RMSSD) formula; 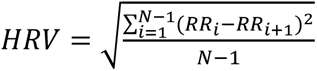 ^69^.

### Heart samples for heart weight to body weight ratio

Mice P70-90 were weighed before they were euthanised via cervical dislocation following deep isoflurane anaesthesia. Their hearts were excised, washed in PBS and blotted dry on paper towel. The individual hearts were then weighed to obtain heart weight to body weight ratio as previously described^70^.

### Proconvulsant assay

Baseline recording was performed for 60 minutes. The mice were then injected intraperitoneally with either vehicle (0.9% sodium chloride) or atenolol (10 mg/kg)^71^. This was followed by a subcutaneous 100 mg/kg PTZ injection 45 minutes later as per previously described^48^. The mice were monitored for a maximum of 60 minutes, after which all surviving mice were culled by cervical dislocation. Cardiac parameters were analysed as above, with ECG analysed at baseline and throughout the entire seizure.

### Statistical analyses

All statistical analyses were performed using GraphPad Prism software (version 9.2.0, Boston, MA, USA). Survival curves were plotted with Kaplan-Meier estimator and analysed using Mantel-Cox log-rank test. All other data were reported as mean ± standard error of the mean (SEM). Data sets were first analysed using a Shapiro-Wilk test for normality. For statistical comparison of two normally distributed data, an unpaired two-tailed Student’s t-test was used. For data where Shapiro-Wilk test returned P < 0.05, a Mann-Whitney U-test was used for two-group comparison. One-way ANOVA with Dunnett’s post-hoc or Kruskal-Wallis test was used for statistical comparison of more than two normally or non-normal distributed data groups respectively. Welch’s correction test was used for groups with unequal variances. Statistical significance was set at P < 0.05. Summary of statistical analyses is listed in Tables 1 and 2.

### Data availability

All requests for materials and resources (data, mouse lines, full-length videos) should be directed to the corresponding author (CAR).

## Acknowledgements

We thank Professor B. London of the University of Iowa for the *Kcnh2*^+/-^ mice and their genetic information; Professor S. Petrou, Associate Professor S. Maljevic, Dr M. Li, Dr T. Karle and Professor I. Forster for technical advice and help with equipment and facility; Dr N. Jancovski for assistance with ECG technique. This work was supported by an anonymous philanthropic gift for SUDEP research (to S.F.B. and I.E.S.). We would also like to acknowledge the CURE foundation, Medical Research Future Fund (#2016012) and National Health and Medical Research Council (#2019804) for support. C.S. and I.E.S. are recipients of National Health and Medical Research Council Practitioner Fellowships (#1154992 to C.S. and #1104831 to I.E.S.) and a Senior Investigator Fellowship (#2016822 to C.S. and #1172897 to I.E.S.). This work was also made possible through the Victorian State Government Operational Infrastructure Support and Australian Government National Health and Medical Research Council Independent Research Institute Infrastructure Support Scheme.

## Author contributions

CAR and MSS conceptualised and designed the study. MSS, AK, ESMS, HML, CEM, AMP and AH performed experiments. MSS, AK, ESMS, HML and CAR performed data analyses. CAR, IES, SFB and MSS acquired funding for the project. MSS and CAR wrote the manuscript. MSS, ESMS, CEM, AMP, IES, CS, SFB and CAR revised the manuscript. All authors approved the final manuscript.

