## Supplemental document for "Atenolol reduces cardiac-mediated mortality in genetic mouse model of sudden unexpected death in epilepsy"

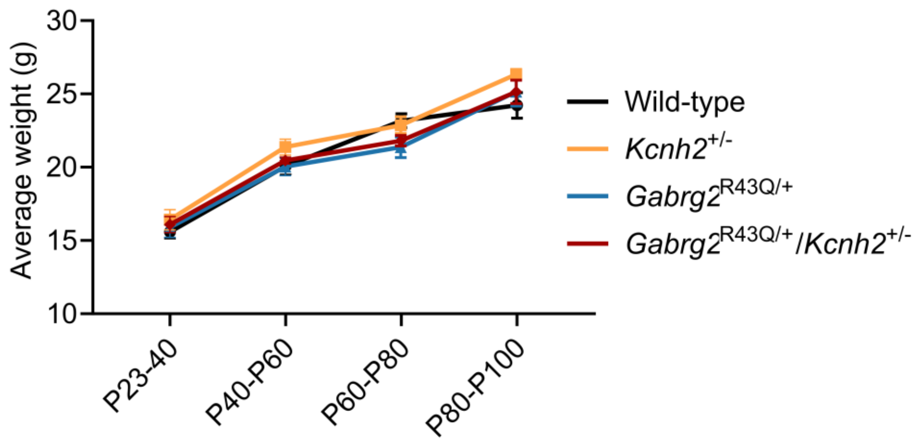


**Supplemental Figure 1: Average weight of littermates of *Gabrg2*$^{\mathbf{R43Q/+}}$ and *Kcnh2*$^{+/-}$ cross.** Littermates from *Gabrg2*$^{R43Q/+}$ and *Kcnh2*$^{+/-}$ cross had similar weights across the four genotypes. Kruskal-Wallis test with Dunn's posthoc at each age group, N=37-45/group, $P > 0.05$.

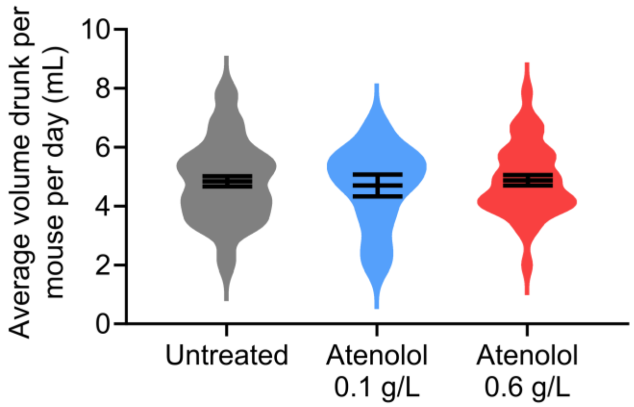


**Supplemental Figure 2: Average volume of water drunk per mouse per day.** Comparison of volume drunk daily per mouse revealed no difference between the groups (Kruskal-Wallis test with Dunn's posthoc, N=9-29 animals per group, $P > 0.05$).

**Supplemental Movie 1: Example video showing a non-terminal spontaneous Racine 4 seizure.**
